# Predicting gene expression from histone marks using chromatin deep learning models depends on histone mark function, regulatory distance and cellular states

**DOI:** 10.1101/2024.03.29.587323

**Authors:** Alan E Murphy, Aydan Askarova, Boris Lenhard, Nathan G Skene, Sarah J Marzi

## Abstract

To understand the complex relationship between histone mark activity and gene expression, recent advances have used *in silico* predictions based on large-scale machine learning models. However, these approaches have omitted key contributing factors like cell state, histone mark function or distal effects, that impact the relationship, limiting their findings. Moreover, downstream use of these models for new biological insight is lacking. Here, we present the most comprehensive study of this relationship to date - investigating seven histone marks, in eleven cell types, across a diverse range of cell states. We used convolutional and attention-based models to predict transcription from histone mark activity at promoters and distal regulatory elements. Our work shows that histone mark function, genomic distance and cellular states collectively influence a histone mark’s relationship with transcription. We found that no individual histone mark is consistently the strongest predictor of gene expression across all genomic and cellular contexts. This highlights the need to consider all three factors when determining the effect of histone mark activity on transcriptional state. Furthermore, we conducted *in silico* histone mark perturbation assays, uncovering functional and disease related loci and highlighting frameworks for the use of chromatin deep learning models to uncover new biological insight.

**Graphical abstract:** 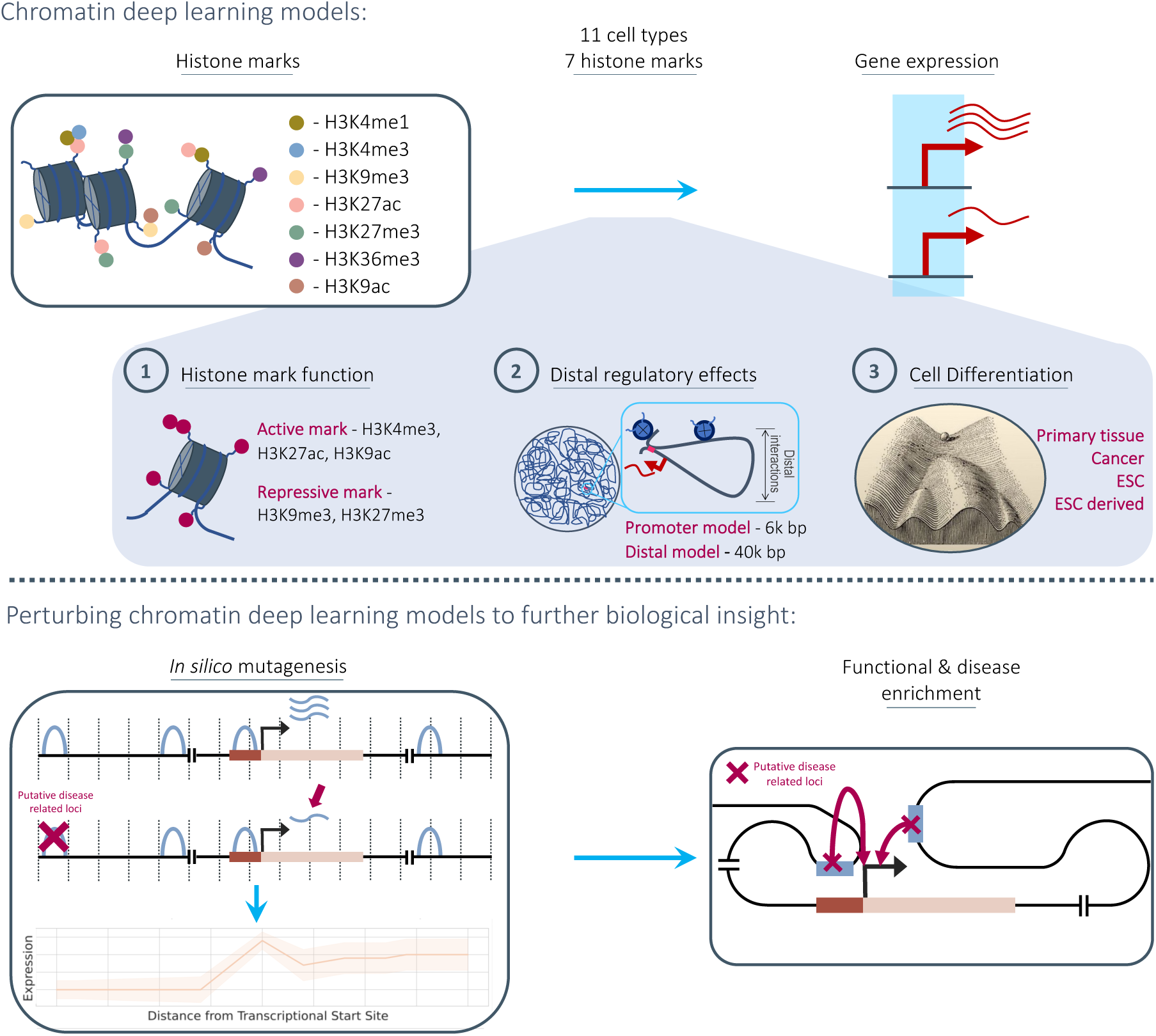

## Introduction

Post-translational modifications on the N-terminal tails of histone proteins, known as histone marks, form a key epigenetic mechanism by which eukaryotic cells regulate transcriptional activity, via altering chromatin structure and interacting with other transcriptional regulators^1,2^. These epigenetic modifications enable cell plasticity without changes to the underlying DNA sequence. Histone mark dynamics in a given cell are mediated by both internal and extracellular queues^3,4^. Alterations in histone modifications have been found to strongly associate with cellular differentiation, cell cycle stages and the development of different diseases^5–8^. For example, as cells mature and differentiate, chromatin accessibility and histone acetylation become progressively restricted throughout their lineage^9^.

Individual regulatory effects of histone marks on transcription have been widely studied: While H3K9ac is associated with active promoter regions^10^, H3K4me1 is found at distal enhancers^11^. However, less emphasis has been placed on the extent to which these histone marks directly regulate gene expression levels. To investigate if transcriptional levels in different cellular contexts can be determined solely from histone modification states, one could correlate levels of histone modifications in regulatory elements with mRNA expression individually. However, this ignores two additional levels of complexity: Firstly, it is well-known that histone modifications interact with other epigenetic factors such as pioneer transcription factors, which in turn have been linked to enhancer activation^9^. Secondly, histone marks themselves act in concert and the interaction between different regulatory elements is not necessarily additive. To circumvent these challenges it is possible to conduct *in silico* experiments, predicting transcriptional levels from histone mark signals where the observed chromatin state is assumed to capture the other contributing epigenetic regulators without directly experimentally observing them.

This method has been applied in previous work, to investigate which histone mark is most predictive of gene expression. For example, addressing this over 20 years ago, Karlić *et al.*^12^ used a linear regression model but only considered histone mark levels at promoter regions, and only tested the effect in a single cell type, CD4+ T-cells. González-Ramírez and colleagues^13^ identified predictive histone marks at promoters and other regulatory regions, leveraging chromatin interaction data from a Hi-C assay to link enhancers to target genes. However, this study also only considered one cell type, mouse embryonic stem cells. Moreover, there is some circularity in the selection of training regions based on derived (from their histone mark data) promoter and enhancer locations and the model’s input measuring the same histone mark levels. Finally, Wang *et al.*^14^, investigated the relationship between histone marks and transcription but inverted the problem, predicting histone mark levels from transcription. For transcription, they used Pro-seq and GRO-seq which labels RNA as it is being transcribed, avoiding issues with RNA degradation^14,15^. The authors used a Support Vector Regression model but again only investigated relationships in the K562 cell line.

Here, we expand on previous research by considering multiple cell types, histone marks and regulatory distances. We investigated the effects of seven histone marks on gene expression (**Table 1**) in eleven human cell or tissue types from the Roadmap Epigenomics Consortium^16^. We will refer to these as cell types hereafter, but note that they also contain tissue samples and cell lines. We used two neural network architectures to predict gene expression: a simple convolutional neural network considering genes promoter regions, and a recently published transformer-based, DNA interaction-aware deep learning architecture called Chromoformer^17^. Chromoformer was originally trained to predict expression using seven histone marks. Here we adapt and retrain the model to predict based on single histone marks, and pairwise combinations of histone marks, to investigate their effect on transcription in isolation. Our work highlights how histone mark function, cellular differentiation and genomic distance to regulatory elements all collectively influence the relationship between histone modification levels and gene expression. We find that there is no universal histone mark which is consistently the most predictive of expression. We recommend that researchers consider all three of these influencing factors when determining the effect of histone mark levels on the transcriptional state of a cell in their work.

**Table. 1.**
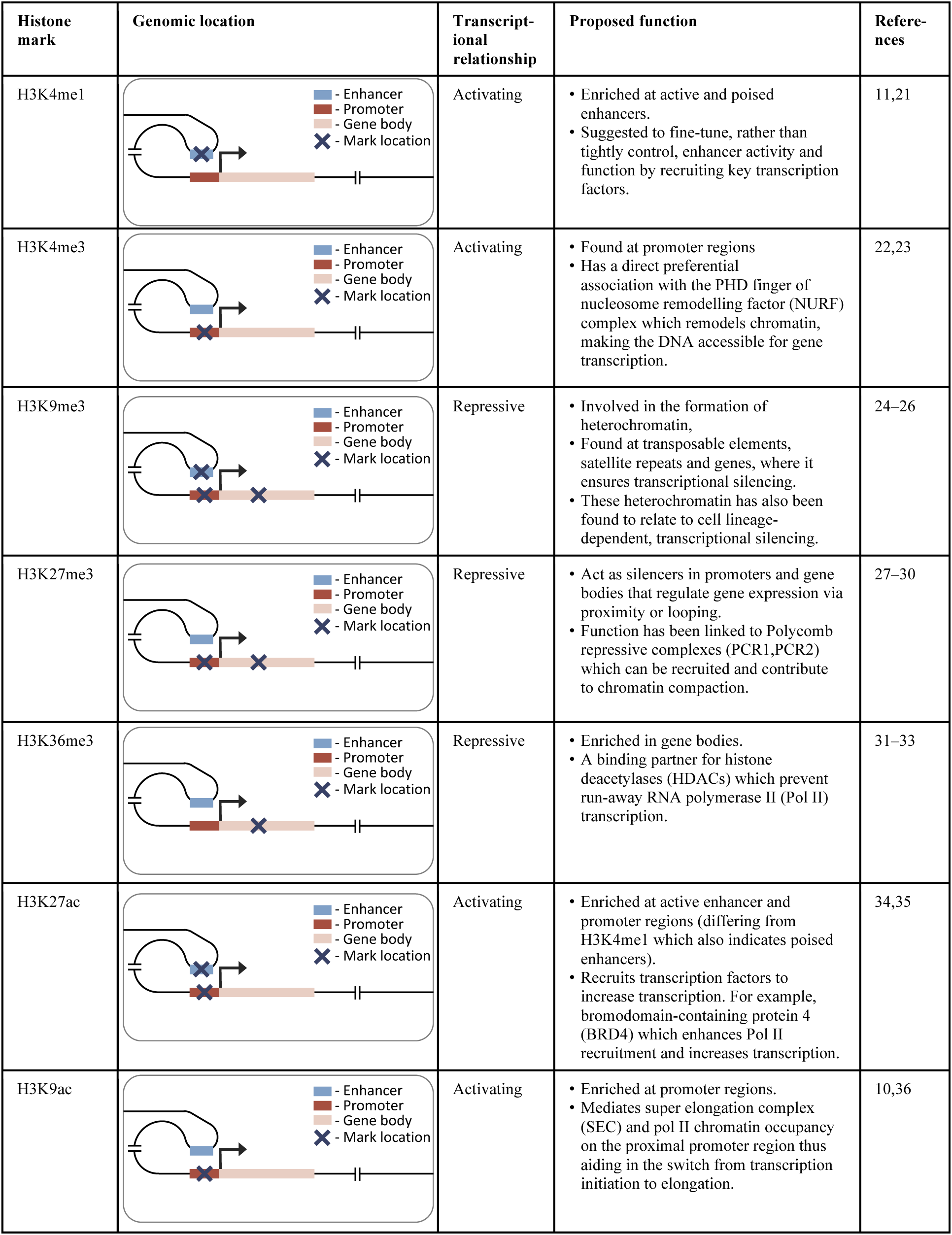
Information on the primary genomic location, transcriptional relationship, and proposed function for the seven histone marks used to predict expression.

In the related field of genomic deep learning where models predict expression or epigenetic marks from DNA sequence, there has been a shift away from arbitrarily benchmarking performance, to prioritising the use of these models to make new biological discoveries^18–20^. This is still lacking for models linking histone mark levels to expression. We aim to address this by outlining a framework to use these models to identify the cell type-specific functional and disease related genomic loci, leading to new biological insights.

## Methods

### Data collection and processing

The data for our analysis was derived from the Roadmap Epigenomics Consortium^16^ and follows the same preprocessing pipeline used by Chromoformer^17^. We used a subset of eleven cell types from Roadmap, for which gene expression, histone mark, and 3D chromatin interactions profiles were available (**Supplementary Table 1**). Specifically we included data from H1 embryonic stem cells (E003), H1 BMP4 derived mesendoderm (E004), H1 BMP4 derived trophoblast (E005), H1 derived mesenchymal stem cells (E006), H1 derived neuronal progenitor cultured cells (E007), HUES64 embryonic stem cells (E016), Liver (E066), Pancreatic islets (E087), A549 EtOH 0.02pct lung carcinoma (E114), GM12878 lymphoblastoid (E116) and HepG2 hepatocellular carcinoma (E118). TagAlign-formatted, ChIP-seq read alignments for seven histone marks - H3K4me1, H3K4me3, H3K9me3, H3K27me3, H3K36me3, H3K27ac, and H3K9ac were used. For consistency, the data was subsampled to 30 million reads and reads themselves were truncated to 36 base-pairs, reducing possible read length biases. The alignments were sorted and indexed using Sambamba^37^ v0.6 and read depths for each base-pair position were derived along the hg19 reference genome using Bedtools^38^ v2.23. Both the promoter and distal models used the averaged log2-transformed 100 base-pair binned signal with our distal model also averaging at 500 and 2,000 base-pairs to also use as model input features. Using three different resolutions of the histone mark signal in the distal model is intended to represent prior knowledge that epigenetic regulation operates on differing scales and has been shown to improve performance of other models^39^. Promoter-capture Hi-C 3D chromatin interaction data^40^ was incorporated into the distal model and mapped to the Roadmap cell types using the same approach as previously described for Chromoformer^17^. Reads Per Kilobase of transcript, per Million mapped reads (RPKM) normalised gene expression levels from protein coding genes were downloaded from Roadmap for the matching cell types. Both the promoter and distal models predict the log2-transformed RPKM (log_2_(RPKM+1)) gene expression levels. The TSS was identified using RefSeq annotations (release v210)^41^ for each gene. Our final dataset included a total of 18,955 genes.

### Promoter model

Our promoter model is a custom convolutional neural network, similar in architecture to DeepChrome^42^. The model takes in a symmetrical 6,000 base-pair genomic window averaged at 100 base-pair bins, centred on the TSS of the gene of interest. The model architecture is composed of three standard convolutional blocks. These blocks each consist of a one-dimensional convolutional layer, batch normalisation, rectified linear unit (ReLU) activation, max-pooling and dropout. This was followed by two fully-connected blocks, which have dropout (in the first block), a dense layer, ReLU activation and a final output layer with linear activation. The convolutional blocks and their sliding window converted the histone mark signal into a position-wise representation highlighting genomic loci that correlate with expression. The fully connected blocks scaled down the size of the representation, to finally produce a single score representing the gene’s RPKM. The size of each layer is provided in our github repository (https://github.com/neurogenomics/chromexpress).

### Distal model

Our distal model architecture was based on the Chromoformer model^17^. This is an attention-based model which uses cell type-specific promoter capture Hi-C data to identify interacting regions in a 40,000 base-pair genomic window centred on the TSS. This approach captures the histone mark signal both at the TSS and at putative cis-regulatory regions. The model has three independent modules at different resolutions (100, 500 and 2,000 base-pairs), producing a multi-scale representation of the histone mark landscape. Each module goes through a transformer block before being combined and passed through a full-connected block with ReLU activation and a final output layer with linear activation. The architecture of the model is discussed in more detail in the original publication^17^.

### Model training

The same model training approach was used for both the promoter and distal model. Model training and evaluation was based on a 4-fold cross-validation regime to give a stronger estimate of model performance. The total 18,955 genes were split into four sets, 5045, 4751, 4605, and 4554 respectively, with genes from the same chromosome in the same split to avoid data leakage^43^. For every fold, one set was used as the blind test set, while the other three sets were used for model training and validation. Performance on the test set for each fold was measured with Pearson’s correlation coefficient. A separate model was trained for each histone mark, cell type and cross-validation fold combination.

The models were trained using the ADAM^44^ optimiser with default parameters with a batch size of 64 over a maximum of 100 epochs. An early stopping regularisation was implemented based on the model’s validation loss with a patience of twelve epochs. The initial learning rate was set at 0.001 and decayed by a factor of 0.2 when the loss did not improve after a patience of three epochs. Mean squared error (MSE) was used as the loss function.

### Histone mark levels

Histone mark occupancy was measured separately for our promoter and distal models and for each cell type, histone mark and cross-validation fold. It was measured as the average log2-transformed, 100 base-pair binned read depth of the histone mark signal. For the promoter model, this signal was taken from the 6,000 base-pairs around the TSS of each gene and for the distal model, from the full 40,000 base-pairs.

### Gene expression state

Histone mark occupancy was measured for both active and inactive genes. A gene was defined as active or inactive based on whether its expression level is above or below the median for that cell type. This approach was first implemented in DeepChrome^42^ and has been frequently used in the literature^17^.

### Correlation analysis

We measured agreement in model performance for both our promoter and distal models by matching the cross-validation fold and cell type for each model trained on a pair of different histone marks. The Pearson correlation coefficient was used to quantify agreement.

### *In silico* perturbation of histone mark activity levels

*In silico* histone mark perturbation was performed on the distal model trained on a single mark. Although we have trained Chromoformer with multiple histone marks as input, we chose to use it trained on a single mark for the perturbation analysis since perturbing one mark will likely affect the histone mark occupancy of other marks in the same region which would not be possible to accurately account for in the model.

We selected one active and one repressive mark which are found at both promoter and distal regulatory elements - H3K27ac and H3K27me3. Perturbation experiments were carried out on active genes for the active mark model and inactive genes for the inactive mark model (see Methods - Gene expression state), to measure the effect on expression of reducing the levels of the histone mark. The predictions from the different k-fold versions of the model were averaged, similar to the approach commonly used in sequence to expression models^45–48^. For the promoter histone signal, the full 6,000 base pairs around the TSS were perturbed, whereas for the distal histone signal, bins of 2,000 base pairs across the 40,000 base pair receptive field were perturbed iteratively (similar to the approach for DNA sequence perturbation used by the genomic deep learning model CREME^49^). The implemented perturbation levels were between 0 and 1 inclusively in 0.1 steps. The code to perform the *in silico* histone mark perturbation is available on our github repository (https://github.com/neurogenomics/chromexpress).

As well as averaging the predictions from the different k-fold versions, we also tested the correlation between the different folds to ensure the model is learning consistent regulatory code. Moreover, we benchmarked this concordance against Borzoi^45^, a genomic deep learning model with the current largest receptive field of 524,000 base pairs. Here, for a fair comparison, we only tested Borzoi’s concordance in the centre 40,000 base-pairs to match Chromoformers receptive field, for the RNA predictions in the same cell types as those used in our analysis and added up to 500 random genetic variants to the sequences of 1,000 genes to match our perturbation test.

### *In silico* perturbation enrichment in quantitative trait loci studies

The averaged *in silico* perturbation experiments on the active (H3K27ac) model for each cell type were filtered to those greater than 6,000 base-pairs upstream of the TSS to concentrate on distal, cell type-specific regulatory regions as opposed to promoter regions or gene bodies (which have a median length of ∼25,000 base pairs^50^, longer than the downstream receptive field of the model). These were next sorted based on their predicted change in expression and split into deciles.

The fine-mapped expression quantitative trait loci (eQTL) data based on the UK Biobank population was sourced from Wang et al., 2021^51^. Causal SNPs were identified from those in linkage disequilibrium (LD) using FINEMAP v1.3.1^52^ and SuSiE v0.8.1^53^. The resulting fine-mapped SNPs were filtered to those with a SuSiE causal probability (posterior inclusion probability (PIP)) > 0.9 in the tissue of interest and with a PIP <0.1 in other tissues to get just the high confidence, tissue-specific fine-mapped SNPs. The ROADMAP cell types were matched to five of the tissues used in eQTL study where the tissue assayed were identical across the two (available on our github repository: https://github.com/neurogenomics/chromexpress). To test for enrichment of the fine-mapped, tissue-specific SNPs, a bootstrap sampling experiment was implemented where the proportion of SNPs found in each decile were compared against 10,000 randomly sampled regions from all deciles. P-values were derived and adjusted using false discovery rate (FDR) correction for multiple testing. Since the distal model uses Hi-C chromatin interaction data as input, we also ran this eQTL enrichment test on the matched cell type and gene, significant promoter capture Hi-C interactions to compare against the model’s enrichment performance. To match the model’s tested regions, the chromatin interaction data was filtered to just those upstream of the gene. Scripts detailing the approach are available on our github repository (https://github.com/neurogenomics/chromexpress).

### *In silico* perturbation disease enrichment

To test for disease enrichment, the top decile of the same averaged *in silico* perturbation experiments on the active model, filtered to just those greater than 6,000 base-pairs upstream of the TSS, were considered. For this analysis, predictions in the liver and neuronal progenitor cells (NPCs) were used due to their potential respective relationships with liver and neuronal diseases. A third group of regions comprising the top decile across all cell types was included to look for cell type-consistent disease enrichment.

Summary statistics for genome-wide association studies (GWAS) for liver diseases - non-alcoholic fatty liver disease (NAFLD)^54^ and hepatitis^55^, glial diseases - Parkinson’s^56^ and Alzheimer’s^57^ and neuronal diseases - amyotrophic lateral sclerosis (ALS)^58^, schizophrenia^59^, autism spectrum disorder^60^ and bipolar disorder^61^ were downloaded from the IEU GWAS portal^62^ and the BioStudies database^63^ and were uniformly processed with MungeSumstats v1.11.3^64^ (default settings, converting build to hg19 where necessary and saving in ‘LDSC’ format).

We applied stratified LD score regression (s-LDSC)^65^ v1.0.1 (https://github.com/bulik/ldsc) to test for disease enrichment. Specifically, annotation files for each of the three groups of genomic loci were first created with Phase 3 of the 1000 genomes reference. Followed by the generation of LD scores with a window size of 1 centiMorgan (cM) i.e. approximately 1 million base pairs, filtering to HapMap3 SNPs to match the baseline model. Finally, the enrichment analysis was run for the GWAS summary statistics across the three groups as well as those in the baseline model whilst excluding the major histocompatibility complex (MHC) (due to the known difficulties predicting LD in this region)^65^.

## Results

### Active histone marks prove most informative at promoter regions

We first focused on the performance of histone mark levels from the promoter regions of the gene of interest using our promoter model. Overall, we found H3K4me3, a mark located in active promoter regions^22^, to be the best performing (**Fig. 1a**). Moreover, active promoter marks H3K4me3, H3K27ac^34^ and H3K9ac^10^ made up the three top performing marks, all with a Pearson’s correlation above 0.73. The fourth best performing histone mark, H3K4me1, is found at active enhancers^11^. It likely performed worse than the other active marks due to the limited range of the model, which only took into consideration a gene’s promoter region. Repressive marks proved less informative with H3K9me3^24^, H3K27me3^27^ and H3K36me3^66^ making up the three worst performing marks. Importantly, there was high variability in performance across the histone marks with a correlation difference of 0.25 between the best active and worst repressive mark (range of Pearson correlation coefficients: 0.52 - 0.76).

**Fig. 1.**
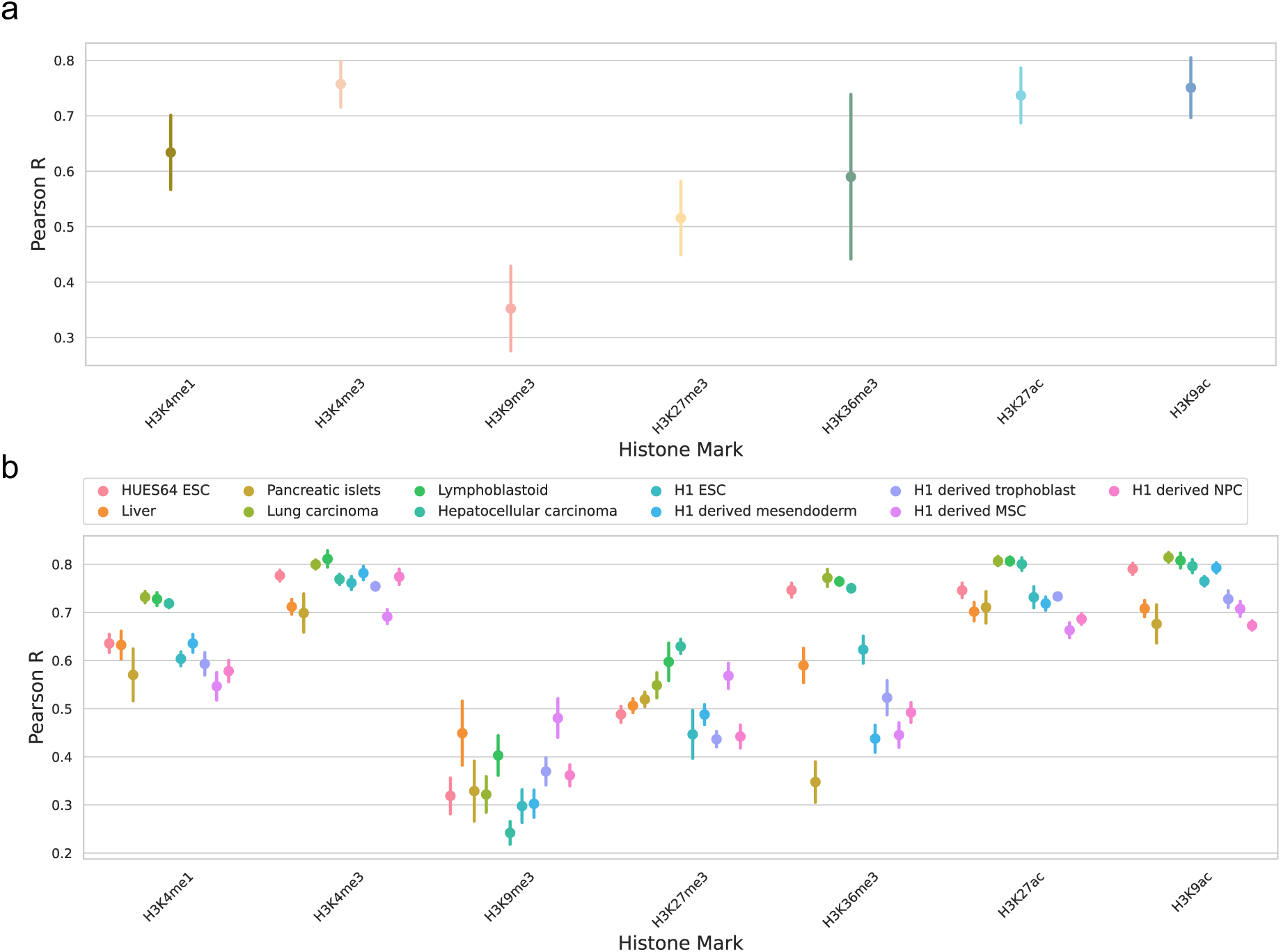
The performance of the promoter model shows the best gene expression predictions based on active histone marks H3K4me3, H3K27ac and H3K9ac. (a) Model performance measured by the Pearson correlation coefficient on the blind test sets across each histone mark. The whiskers represent the standard deviation across the different cell types and the 4-fold cross-validation. (b) Performance split by different cell types from Roadmap.

### The predictive performance of active and repressive marks differ based on cell state at promoter regions

Splitting the models’ performance across the different cell types highlighted histone mark groups with similar variations in their scores across cell types (**Fig. 1b**). This is most notable for active promoter marks (H3K4me3, H3K9ac and H3K27ac). To formally evaluate this trend, we calculated the correlation between the models’ predictions across the different genes, cell types, histone marks and cross-validation folds (**Fig. 2a**). One distinct group of highly correlated histone marks were apparent (highlighted in blue in **Fig. 2a**), corresponding to the active histone marks previously observed. Interestingly, H3K9me3 showed the lowest correlation with the other histone marks, including with H3K36me3 and H3K27me3, the other repressive marks. The samples collected for Roadmap^16^ can be classified into those taken from Embryonic stem cells (ESC), cells differentiated from ESCs, adult bulk tissues and cancer cell lines (**Supplementary Table 1**). Active histone mark activity levels were significantly more predictive of expression in ESC than primary tissue whereas the opposite was noted for repressive histone mark activity levels which was more predictive in primary tissues than ESC (**Fig. 2b**). The multi-modal performance, visible in **Fig. 2b**, is the result of a combination of the histone mark and cell type being tested (**Supplementary Fig. 1**). These results indicated that the most accurate method by which to predict gene expression from the promoter region depends on the extent to which the cell type of interest has differentiated - active marks like H3K4me3, H3K9ac and H3K27ac were most predictive for cells at earlier stages of their differentiation process whereas repressive marks, like H3K9me3, fared better in fully differentiated tissues or cells.

**Fig. 2.**
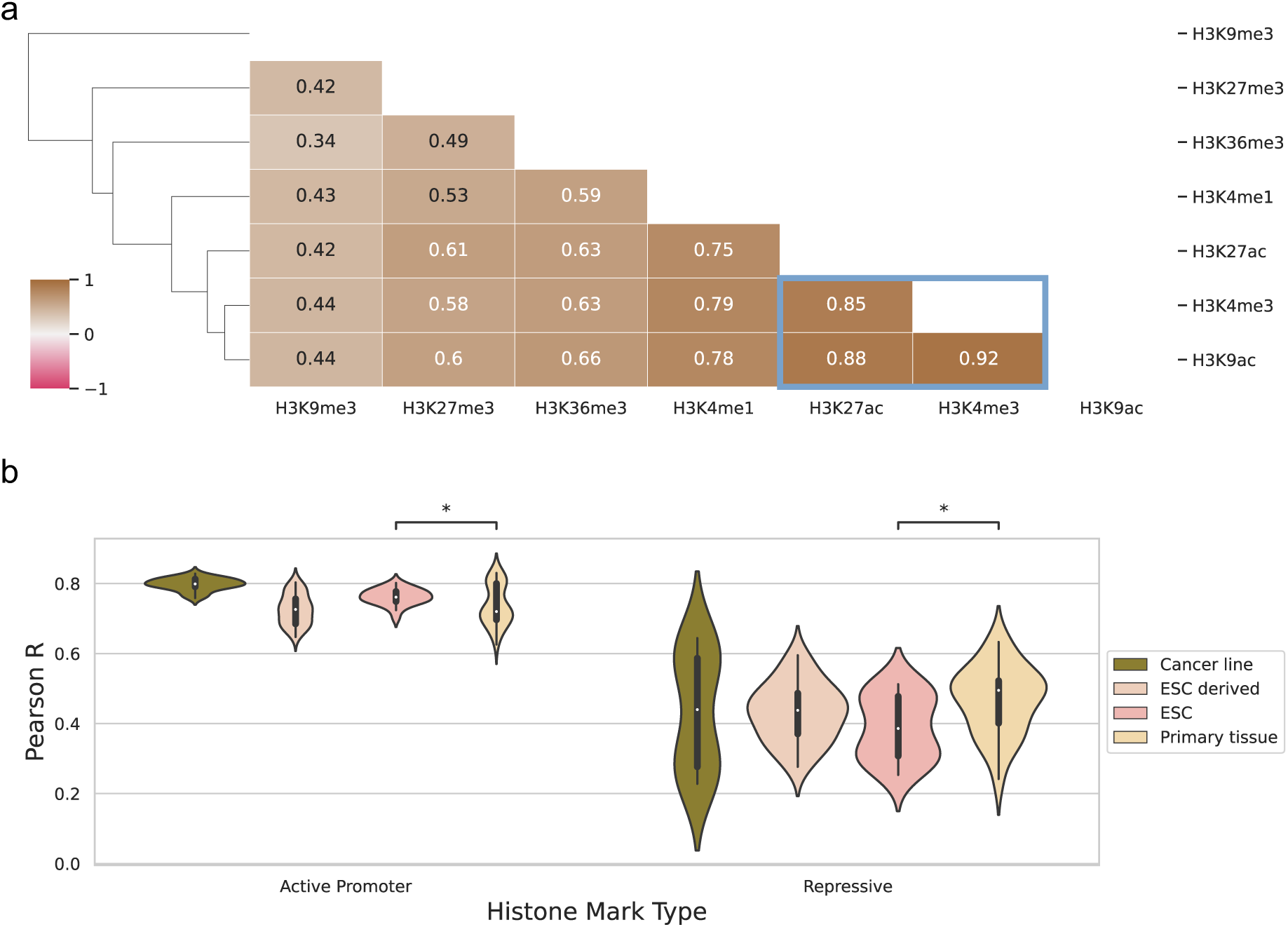
The predictive performance of the promoter model across samples clusters by histone mark function. (a) Correlation matrix of the promoter model’s predictions by the different histone marks across each gene, cell type and cross-validation fold. H3K4me3, H3K9ac and H3K27ac were characterised by high positive pairwise correlations (highlighted in blue). Bars along the y-axis show the hierarchical clustering dendrogram. (b) Violin plot of the model’s performance on active and repressive histone marks measured by their Pearson correlation coefficient on the blind test sets. The cell type performance is grouped by the cell state - Embryonic stem cell (ESC), ESC derived cell, adult primary tissue or cancer cell line. Significance was based on the Mann-Whitney U-Test with false discovery rate (FDR) correction for multiple testing, with p-value indicator: * < 0.05.

### Higher histone mark levels result in better predictive performance at promoter regions

To investigate what was driving the difference in performance for active and repressive histone marks in the different cell states, we measured the histone mark levels in the promoter regions for highly and lowly expressed genes (**Fig. 3**, see **Methods** for details). We observed a strong correlation between model performance and histone mark levels for active histone marks in highly expressed genes, and conversely, for repressive marks in lowly expressed genes, indicating that the model learns the functional significance of the different histone marks (**Fig. 3a**). For highly expressed genes (**Fig. 3b**), repressive marks, H3K9me3, H3K36me3 and H3K27me3, had higher histone mark levels in ESCs than primary tissue (although for H3K9me3 this was not significant after multiple test correction). Conversely, active marks, H3K4me3, H3K9ac and H3K27ac, showed varying activity across primary tissue and ESCs. For lowly expressed genes (**Fig. 3c**), histone mark levels tended to be higher in ESCs than in primary tissue across histone marks. Overall, our analysis highlighted that higher histone mark levels in a gene where the expression status matched the function of the histone mark (active vs repressive), led to better performance of the model.

**Fig. 3.**
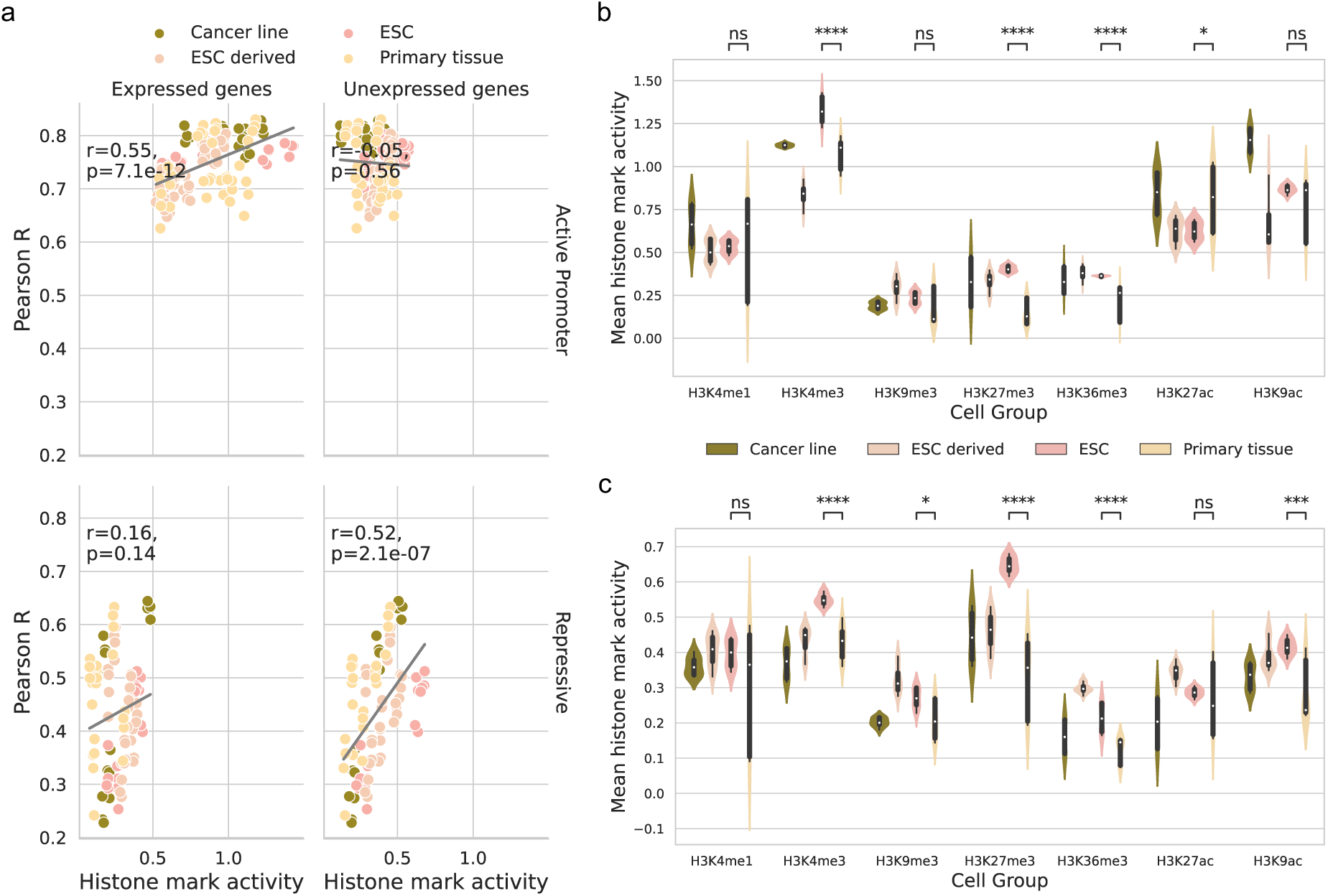
Higher histone mark levels are associated with better predictive performance. (a) Correlation between the histone mark levels and model performance for each cell type, k-fold combination. This is split by repressive and active marks, as well as highly and lowly expressed genes. (b) Violin plot of histone mark levels measured by the average log2-transformed, read depth in the promoter region (6,000 base-pairs around the transcription start site) for highly expressed genes. The cell types are grouped by the cell state - Embryonic stem cell (ESC), ESC derived cell, adult primary tissue or cancer cell line and averaged at the level of cell type and k-fold. (c) Average histone mark levels for lowly expressed genes. Significance was based on the false discovery rate (FDR) multiple test correction, with p-value indicators: **** < 1e-4, *** < 1e-3, ** < 1e-2, * < 0.05, ns >= 0.05.

### Active marks are most predictive in the distal model

To consider histone mark levels outside of the genes’ promoter regions, we next tested a model architecture with a much larger receptive field (up to 40,000 base-pairs around the TSS). We used Chromoformer^17^, a transformer-based architecture, which accounts for distal histone mark levels, weighting important genomic regions using cell type-specific DNA interaction data. We trained this distal model on each single histone mark and benchmarked the performance across histone marks, and also against the performance when trained on all seven histone marks combined (**Fig. 4a**). Again, we found H3K4me3, H3K9ac and H3K27ac to be the top three performing marks with very little difference in overall performance between them (mean Pearson R of 0.762, 0.757 and 0.749 respectively). All three histone modifications are active marks while only H3K27ac^34^ is found at both promoters and distal regulatory regions (enhancers).

**Fig. 4.**
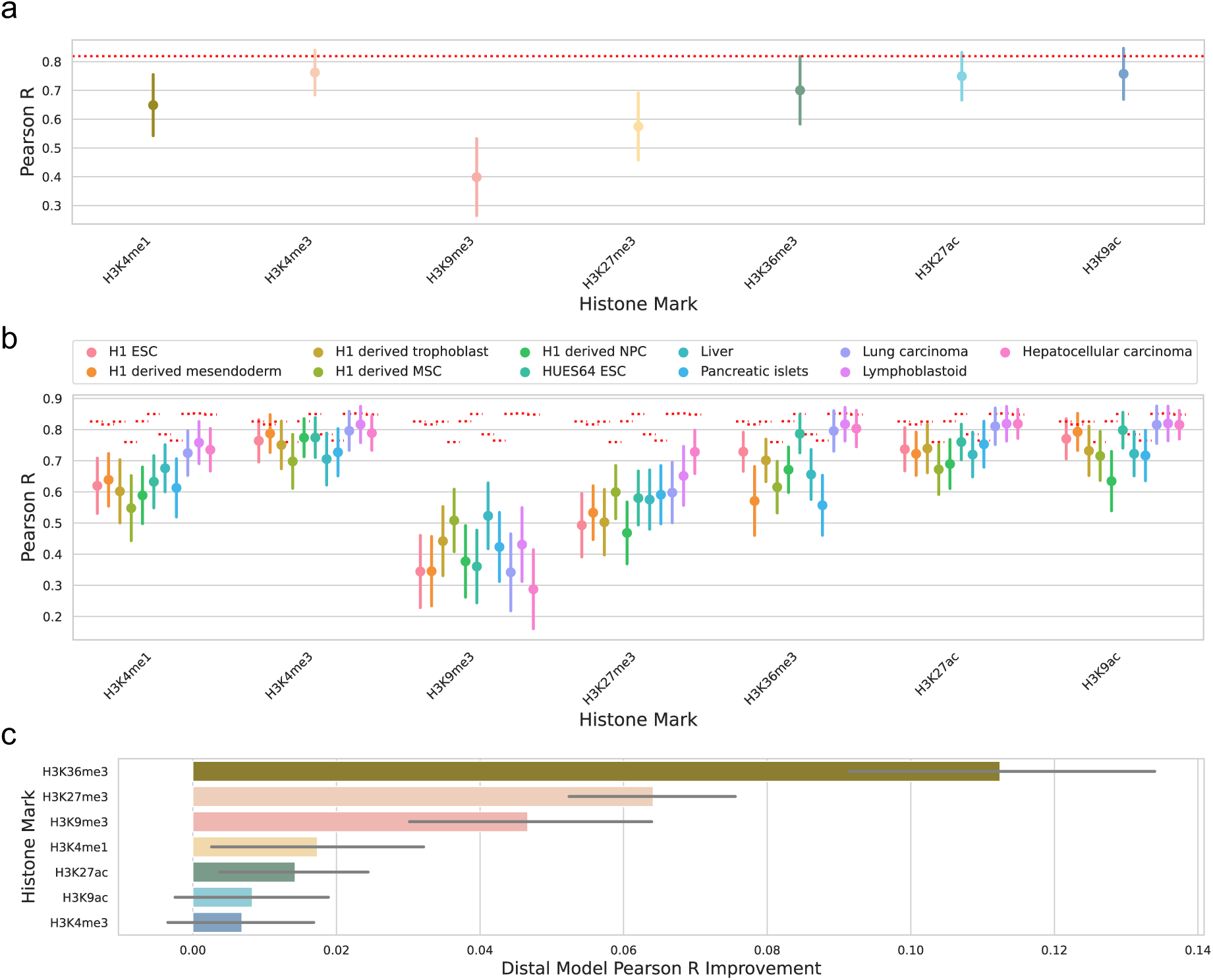
Active histone modifications performed best at predicting gene expression in the distal model. (a) Model performance was measured by the Pearson correlation coefficient on the blind test sets for each histone mark. The whiskers represent the standard deviation across the different cell types and the 4-fold cross-validation. The red dashed line shows the model’s performance when trained on all seven histone marks together. (b) Predictive performance is shown split by the different cell types in Roadmap. (c) The improvement in performance for each histone mark by expanding the receptive field outside of the promoter region with the distal model. The largest increase in performance is observed for H3K36me3. The whiskers indicate the standard deviation across cell types and folds.

### Receptive field expansion and multi-histone mark predictions yield diminishing returns

The performance of incorporating distal histone mark levels in a model consistently but marginally increased the Pearson correlation coefficients. The range of improvement in correlation varied from 0.01 - 0.15, despite the substantial increase in receptive field and model architecture complexity (**Fig. 4c**). Compared to the promoter model, although the same histone mark performance ranking was observed, marks which are known to affect genomic locations outside of promoter regions showed the highest relative improvement, specifically H3K36me3, H3K27me3 and H3K27ac (**Fig 4c**, **Table 1**).

The relationship between histone mark type and cell state found for the promoter model, where active histone marks were more predictive in ESC and repressive in primary tissues, was similarly observed for the distal model (**Fig. 4b, Supplementary Fig. 2a-b**). Moreover, we investigated the histone mark levels across the full 40,000 base-pair receptive field of the distal model and found the same trend as for the promoter model where the model picks up on known biology of histone mark prevalence: We observed strong correlations between model performance and histone mark levels for active histone marks around highly expressed genes, and conversely, for repressive marks around lowly expressed genes (**Supplementary Fig. 2c**). To further interrogate the contributions of histone marks to the prediction of expression in our distal model, we benchmarked performance across pairs of histone marks with the top three performing marks, H3K4me3, H3K9ac and H3K27ac (**Fig. 5a**). All combinations with the top three histone marks were better performing than any of the top three marks by themselves. However, this improvement was relatively small (< 0.04 mean increase in Pearson R for the best histone mark combination) and was only significant for a handful of combinations.

**Fig. 5.**
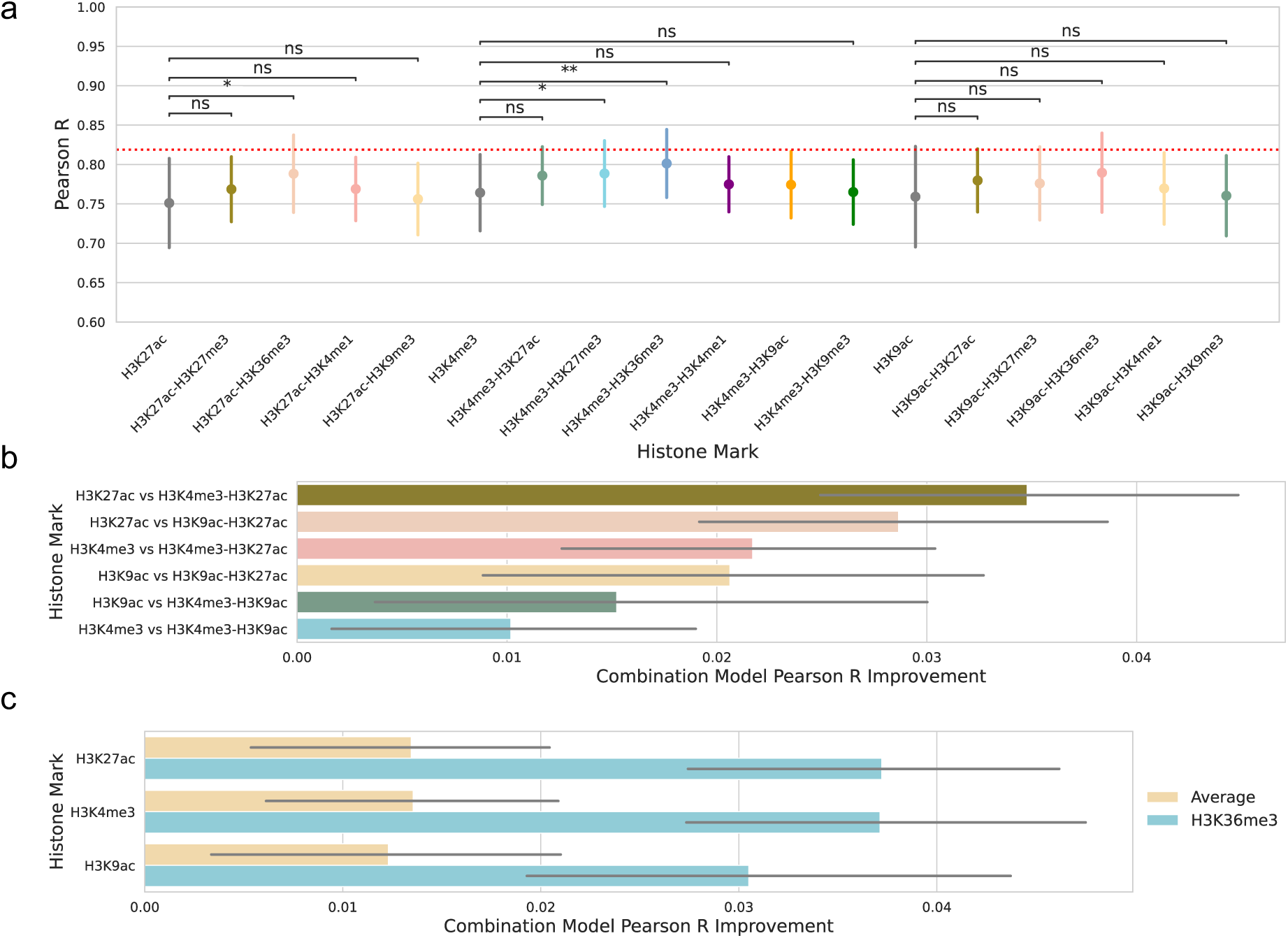
Pairwise combinatorial performance of distal model in predicting gene expression based on two histone marks. (a) Performance was measured by the Pearson correlation coefficient on the blind test sets across combinations of histone marks. The whiskers represent the standard deviation across the different cell types and the 4-fold cross-validation. Data is averaged at the level of cell type and k-fold. The red dashed line shows the model’s performance when trained on all seven histone marks together. Significance based on false discovery rate (FDR) multiple test correction, with p-value indicators: ** < 1e-2, * < 0.05, ns >= 0.05. (b) Performance improvement for combinations of the top three performing marks from the single histone mark distal model. (c) Performance improvement over the single histone mark distal model for combinations using H3K36me3 versus the average of the other marks.

This result highlights that incorporating additional histone marks in the prediction led to a consistent, albeit small, boost in performance, regardless of the histone mark type (active or repressive). This became most evident when we combined pairs of the top three performing marks: All three are active marks and two are confined to promoter regions, but their combination still resulted in improved performance over the individual marks (**Fig. 5b**). Notably though, combinations including H3K27ac, an enhancer mark, showed greater improvement over combinations including only promoter marks. Importantly, combining the top three marks with another mark showed a mean improvement of 0.043, compared to a mean improvement of 0,062 when using all marks (red dashed line in **Fig 5a**). This means that adding an additional five histone marks to the distal model would increase performance (Pearson R) by another 0.02, highlighting the diminishing returns of including additional histone mark information.

Our analysis further highlighted the combinatorial predictive capabilities of H3K36me3 (**Fig. 5a,c**). H3K36me3 is a repressive mark^67^ with strong distal effects on gene expression. This mark showed the largest improvement from the promoter to the distal model (**Fig. 4c**) and when combined with the top three scoring histone marks, was its best performing pair, even when compared to combinations of the top three performing marks (**Fig. 5a**) and far improved performance compared to the addition of the other marks (**Fig. 5c**). Conversely, when paired with the repressive mark H3K27me3, H3K36me3 did not result in the best performing pair (**Supplementary Fig. 3**) but did still improve performance on the individual mark. This highlights the complementary information H3K36me3 provides in addition to the top three performing active promoter and enhancer marks.

We also investigated whether the performance increase for the top three marks when coupled with H3K36me3 was driven by bivalent genes. Bivalent genes are characterised by the presence of both repressive and active histone mark signals at their promoter and are known to silence developmental genes in ESCs while keeping them poised for activation^68^. However, the model performance in bivalent genes for ESCs did not notably improve over non-bivalent genes (**Supplementary Fig. 4**).

### *In silico* histone mark perturbation prioritises functionally relevant genomic loci and disease relevant cell types

Up to this point, our work has shown the performance of chromatin to expression deep learning models and how they encode known biological relationships between histone mark levels and expression. However, none of this highlights new information about gene regulation or disease. *In silico* perturbation enables experimentation to investigate the effect on gene expression of varying histone mark levels in a cell type and gene-specific manner that would be impractical to undertake experimentally *in vitro* or *in vivo*.

We first investigated the effect of *in silico* perturbation experiments at an aggregate level, varying levels of histone mark activity, as well as distances from the TSS using the distal model trained on one active (H3K27ac) and one repressive (H3K27me3) mark (**Fig. 6**). We used models trained on single marks to avoid issues where perturbing one type of histone mark will affect another mark’s activity in the region. Here, we permuted either the entire TSS or the distal regions in 2,000 base pair bins (see **Methods** - *In silico* perturbation of histone mark activity levels). Furthermore, we averaged predictions across the four k-fold model versions, a standard approach *in silico* mutagenesis experiments for genomic deep learning models^45,46,48^. This step may not have been required given the notably high correlation between the different models’ predictions (**Supplementary Fig. 5a**), which was on par with, if not slightly better, than that of the genomic deep learning model Borzoi^45^ (**Supplementary Fig. 5b**). This indicates that models trained on histone mark levels show a similar ability to learn consistent regulatory code across differing training sets compared to that of their genomic counterparts. For the active model, we noted a clear relationship between reducing histone mark levels and large predicted decreases in expression at the promoter (**Fig. 6a**) while we found only minor decreases at distal regulatory regions (**Fig. 6b**). This matches the known functional relationship between promoter and enhancer activity with expression and also the *in silico* perturbations of DNA sequence in genomic deep learning models^46^. However, reducing repressive histone mark levels showed little relationship with increased expression (**Fig. 6c-d**) which may be due to the repressive mark’s relatively worse performance overall (**Fig. 4a**). The relationship between perturbation and distance and their effect on expression is more apparent when we view the predicted quantile change in expression by distance from the TSS while removing histone mark activity completely (**Fig. 6e-f**). Here, we saw the highest predicted change near the TSS, reducing as distance to the TSS increases for both the active and repressive model. Interestingly, this reduction was not symmetrical upstream and downstream of the TSS, with downstream loci having a greater effect on expression on average. We believe this was a result of the length of the gene body (median length ∼25,000 base pairs^50^), which would incorporate the entire downstream receptive field of the model for most genes and thus lend to greater importance for RNA-seq predictions rather than assays of transcription initiation like CAGE-Seq^69^. Our analysis highlighted that on average, perturbations to histone mark signals in the gene body had a greater predicted effect than distal regulatory regions upstream.

**Fig. 6.**
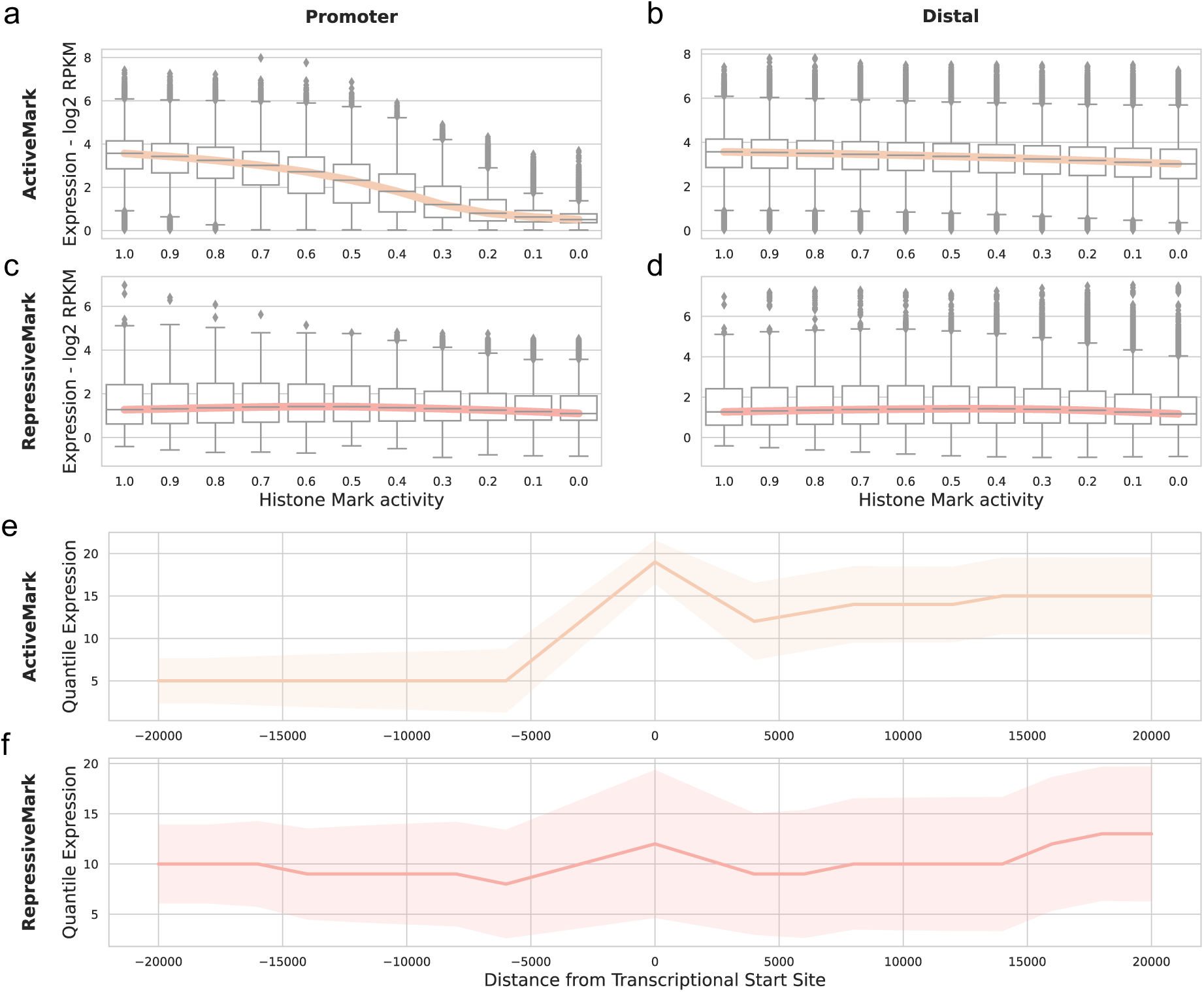
Effect of *in silico* histone mark perturbation on expression. The effect on predicted expression (y-axis) of changing the proportional levels of measured histone mark activity (x-axis) for all cell types and genes, averaged across the four k-fold models. The distal model trained on a single histone mark was used to measure the effects of a perturbed active mark - H3K27ac (a,b) or a repressive mark - H3K27me3 (c,d) in 2,000 base pair bins for distal or at the promoter. (e,f) The effect of distance on expression change is shown when the histone mark activity is completely removed at a specific location for the active (e) or repressive (f) mark. The distribution of all gene expression changes in all cell types split into 20 quantile bins.

We next considered whether these *in silico* perturbation experiments could be used to gain insight into the cell type-specific regulatory function in gene expression and disease, using genetic variants to test for functional and disease enrichment (**Fig. 7**). Given that the active model captured known biological relationships in the *in silico* perturbation, we focused on this model’s perturbation experiments. Moreover, we only considered upstream predictions to capture distal regulatory regions which vary across cell types as opposed to promoter and gene body signals.

**Fig. 7.**
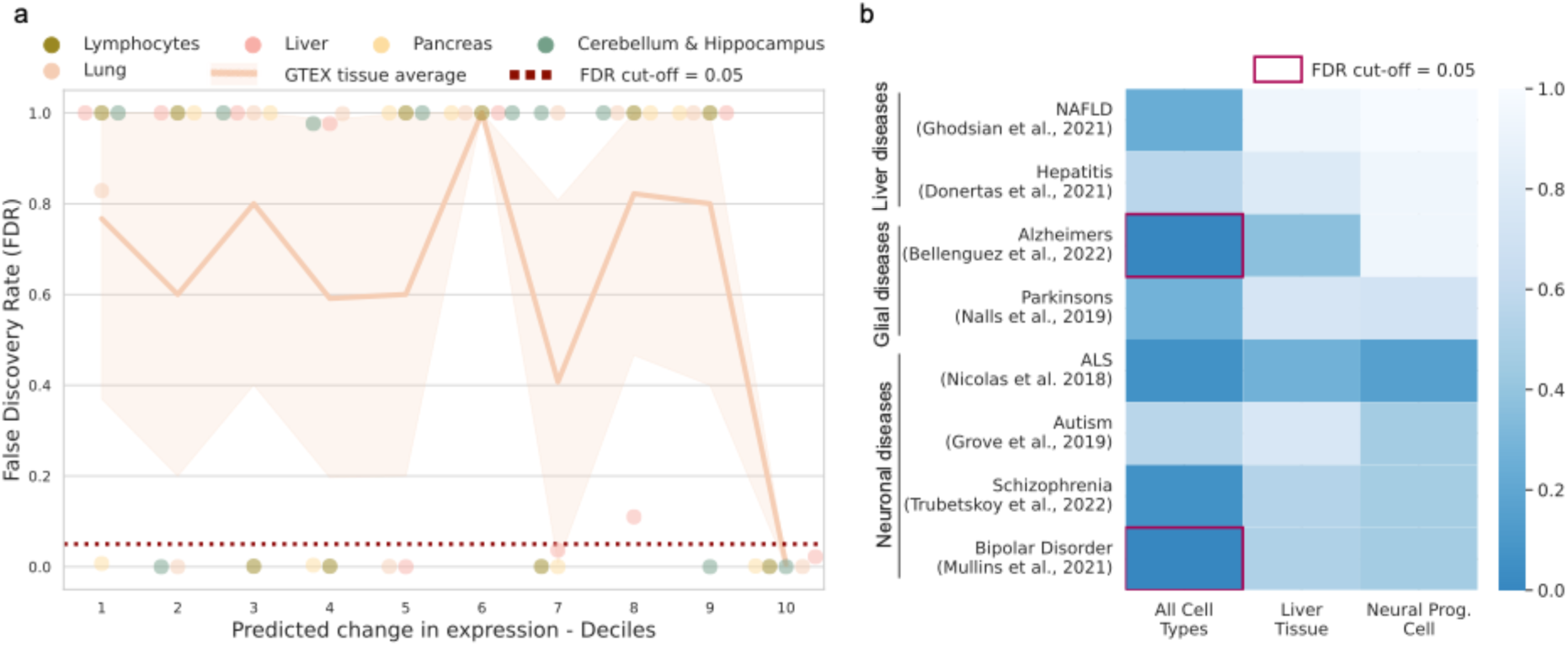
***In silico* histone mark perturbation experiment highlights functional and disease enrichment.** (a) Upstream *in silico* histone mark perturbation experiments from the active model were sorted into deciles based on their predicted change in expression (x-axis). Each decile was tested for enrichment of fine-mapped eQTL interactions in matched cell types and compared against bootstrap sampling random upstream loci 10,000 times to generate p-values of enrichment (y-axis). (b) False discovery rate (FDR) P-value enrichment scores for the top decile (based on their predicted change in expression) derived from all cell types (non-cell type-specific), liver tissue or neuronal progenitor cells (x-axis). Enrichment tests were conducted with s-LDSC and genetic variants relating to different diseases (y-axis). FDR adjustment was applied for each GWAS included.

To test the model’s ability to capture functional loci we used a large scale, tissue-specific, fine-mapped eQTL set based on the UK Biobank population^70^. First, we split the loci into deciles sorted based on the model’s predicted change in expression (decile 10 having the largest predicted effect). We implemented a bootstrap sampling test to compare each decile to randomly sampled upstream loci for enrichment of the fine-mapped eQTLs (see **Methods** - *In silico* perturbation enrichment in quantitative trait loci studies). We found significant enrichment for the top decile for all cell types tested (**Fig. 7a**), indicating that the model correctly predicted the loci which contributed most to the cell type-specific gene expression. Furthermore, we wanted to test how much the distal model improves upon the Hi-C chromatin interaction data alone for fine-mapped eQTL enrichment in these same loci, and found an improvement in four of the five tissues (**Supplementary Fig. 6**).

We next tested whether these loci also harbour known disease related genetic variants. We used s-LDSC^65^ with GWAS for eight different liver and brain diseases^54–61^. Test regions were based on the top cell type-specific decile with the largest predicted effect for liver and NPCs, as well as the averaged top decile across all cell types of upstream loci (**Fig. 7b**). S-LDSC measures enrichment of disease genetic variants accounting for the obscuring nature of LD^65^. Here, we used the neuronal cells and liver tissue for their predicted relationship with brain and liver diseases, respectively. We used the top decile for all cell types to look for non-cell type-specific disease enrichment. While none of the expected cell type-specific significant enrichments were detected, the neuronal disease GWAS were tending towards significance in neuronal disease related GWAS. Moreover, two significant associations (Alzheimer’s disease and bipolar disorder) were detected after multiple testing correction for the non-cell type-specific disease enrichment, highlighting the functional importance to disease of regions where changes in the histone mark activity levels was predicted to have a large effect on expression across cell types.

## Discussion

We present the most comprehensive deep learning study of the relationship between histone mark levels and transcription to date. We considered multiple cell types and histone marks at differing receptive fields and demonstrated how the prediction of gene expression is dependent on three key contributing factors - histone mark function, regulatory distance and cellular states.

For our analysis, we used the Roadmap^16^ data repository, benefiting from the standardised experimental approach. We investigated the genome-wide activity of seven histone marks across eleven cell types. For the histone mark ChIP-Seq read alignments, we subsampled to 30 million reads to enable a fair comparison across marks. While for gene expression, we utilised RPKM values which measures the mRNA abundance of transcripts normalised by gene length, avoiding any potential within sample bias. Since model predictions were made in the same cell type as training, there was no need to standardise gene expression levels across cells^71^. To ensure robust benchmarking results, we repeated training of both our promoter and distal models across a 4-fold cross-validation, ensuring the test set genes were grouped by chromosomes to avoid any data leakage^43^ - where data in the training set is related to data in the test set, artificially inflating model performance. To run each cell type, histone mark or combination of histone marks, for each cross-validation fold using both the promoter and distal models, was a computationally intensive task. This resulted in 1,276 training and prediction iterations which were all run using an A100 80GB GPU. To avoid overfitting over such a substantial number of iterations, we automated hyperparameter tuning for both models using a learning rate decay and early stopping regime, holding out an independent validation set of genes for monitoring.

Both our promoter and distal models were developed as quantitative, regression models, predicting a gene’s log_2_ RPKMs, which has been shown to yield better generalisation and interpretability than binary classification models^72^. For the promoter model, following the same approach as past benchmarking work^72^, we implemented an intentionally simple convolutional neural network architecture based a relatively small receptive field around the TSS to compare histone mark performance. On the other hand, our distal model, Chromoformer^17^, was a transformer-based architecture accounting for distal histone mark levels in a weighted manner based on DNA interaction data. One limitation is the receptive field of our distal model which extends 40,000 base-pairs around the TSS. Although this is a large window and computationally intensive to include in such a model, it is a fraction of the known distance at which DNA interactions can occur. For example, Hi-C experiments capture cis-interactions up to 1 million base-pairs away^73^.

The results of our promoter region analysis showed that the active marks H3K4me3 and H3K9ac were the most predictive of gene expression (**Fig. 1**). However, their optimal performance was dependent on the cell state, performing better in ESCs whereas repressive marks like H3K9me3 performed relatively better in adult primary tissues (**Fig. 2b**). We concluded that the stage of cell differentiation was the driving factor of performance for active and repressive marks: Active marks performed better at early stages of lineage commitment whereas repressive marks were more predictive in fully differentiated cells. Furthermore, we noted that for active genes, the relationship between a histone mark’s levels and performance replicated known biology: Observing a strong correlation for active histone marks in highly expressed genes, and conversely for repressive marks in lowly expressed genes (**Fig. 3a**). This highlighted that higher histone mark levels were beneficial for the model, leading to greater predictive performance in the correct context. In relation to histone mark levels at the TSS, we also noted that for inactive genes, histone mark activity is reduced with cell differentiation (**Fig. 3b**). The concept that cell lineage commitment leads to globally lower histone mark levels has been previously noted^74^.

For our distal model, active marks H3K4me3, H3K9ac and H3K27ac were the best performing (**Fig. 4a**). Interestingly, these three marks, two of which are linked to the TSS of genes they regulate, outperformed H3K4me1 which is associated with distal enhancers^11^. One possible explanation for this was investigated in a recent study^46^ which found that, when predicting expression from DNA sequence, a similar attention model prioritised sequences at the promoter region over distal regulatory regions. The reason being that given the multiple choice of enhancer and other regulatory regions and their relatively small influence on gene expression, a model will prioritise the information at the promoter region. We believe the same effect could have contributed to our results whereby the active promoter mark information contributed to expression to a greater degree than distal regulatory regions. This same relationship was clearly notable in our *in silico* perturbation experiments (**Fig. 6e-f**). Moreover, given that H3K4me1 is indicative of poised rather than active enhancers^11^, it would presumably be less predictive of gene activity.

A key point of our findings is the marginal return in performances by: 1. Extending from an intentionally simple local promoter model to an attention based, computationally complex model which accounts for distal histone mark levels (**Fig 4c**), and 2. Increasing the number of histone marks included in the model (**Fig 5b**). We noted that understanding the cell state (undifferentiated or fully differentiated) and the gene state (highly or lowly expressed) of interest and choosing the most appropriate mark for these had a greater impact on performance than the number or receptive field of histone mark levels considered.

Comparing performance across the promoter and distal models, H3K27ac showed the biggest gain of the top three performing marks (**Fig 4**). This was expected given its relationship with active promoters and enhancer regions^34^. However, its performance based solely in promoter regions was still relatively strong, which was reassuring given the mark’s prevalence in complex disease research.

We also trained our distal model on pairs of histone marks, showing that the added performance of incorporating additional histone marks diminishes markedly after this point. This result could benefit researchers wishing to capture transcription in a cell type of interest from limited histone mark information. The paired analysis also highlighted the strong combinatorial performance of H3K36me3. This repressive mark was the best performing choice as a pair with any of our three top marks (**Fig. 5a**) and was the fourth best performing mark of the single histone mark analysis (**Fig. 4a**). H3K36me3 is a canonical mark of transcription, serving as a binding partner for histone deacetylases (HDACs) which prevent run-away transcription of RNA polymerase II^32^. H3K36me3 is generally enriched in gene bodies of mRNAs^31^, outside of the TSS, which may explain its relatively poor performance in the promoter model and why the improvement with the distal model for this mark was the highest of any mark tested (**Fig. 4c**).

Finally, we performed an array of *in silico* histone mark level perturbation experiments, showing the relationship between distance from the TSS and a regulatory region’s effect on gene expression (**Fig. 6**). Our analysis highlighted the very high correlation for the *in silico* perturbations between the different cross-validation fold models (**Supplementary Fig. 5**). A possible advantage of histone mark deep learning models over genomic deep learning models trained on DNA sequence is that they offer an alternative to making single base pair level changes such as genetic variants which are notoriously difficult to interpret, particularly as the genomic window considered by the model increases^45,48,49^. By identifying genomic loci of interest based on perturbing histone mark levels, our model avoids the majority of these issues. Furthermore, using these identified genomic loci, we developed a framework by which such models can be applied to test for both functional and disease enrichment in a cell type-specific manner (**Fig. 7**). The results for the disease enrichment did not recapitulate projected relationships for the neuronal progenitor cells or liver, which could be a result of the imperfect cell type matching, the limited overlap between the disease related genetic variants and the relatively small window of upstream genomic loci considered, or the observed differences in the measured genetic effects on gene expression versus complex traits^75^. This highlights that further work on such models is needed, hopefully increasing the receptive field and using known affected cell types, to capture an association with complex diseases. Importantly, a substantial overlap and large genomic coverage of the loci considered are key recommendations for s-LDSC analyses^65^. This issue when capturing complex phenotypic enrichment is not unique to these models and is also a challenge with genomic deep learning models as highlighted recently^47^. Non-cell type specific genomic loci predictive of gene expression were enriched for GWAS signal for both Alzheimer’s disease and bipolar disorder, indicating the importance of cell type consistent regulatory regions in complex disease. Importantly, this analysis highlights the significance of these predicted genomic loci not only in functional genomics studies (**Fig. 7a**) but also in disease (**Fig. 7b**). Past approaches such as ICEBERG, a pipeline that uses CUT&RUN replicates to create a combined profile of binding events for H3K4me3, have been previously used to uncover functionally relevant regulatory events^76^. However, our approach shows, for the first time, how chromatin deep learning models can be perturbed to uncover genome-wide and cell type-specific functionally and disease relevant regulatory regions.

Overall, our study shows that multiple factors influence the performance of histone marks when predicting gene expression, something which had not explicitly been considered by previous work^12–14^. Our findings suggest that if one wishes to investigate the TSS of genes, promoter-specific active marks H3K4me3 and H3K9ac are the best options. Beyond the promoter region, active marks H3K4me3, H3K9ac and H3K27ac are most predictive, especially in combination with the transcriptionally-associated mark H3K36me3. However, it is worth considering the cell state (differentiated or early stages of lineage commitment) and the state of the genes or interest (highly or lowly expressed) to have a better understanding of the optimally predictive histone marks. Importantly, more effort should be placed on using these models to uncover new biological insights, particularly for phenotypic and disease-based studies. Similar to genomic deep learning models, chromatin deep learning models are capable of capturing functionally relevant genomic loci.

## Data and code availability

The Histone mark ChIP-seq read alignments, RPKM gene expression profiles were downloaded from the Roadmap Epigenomics Web Portal (https://egg2.wustl.edu/roadmap/web_portal/index.html). The promoter capture Hi-C experiments were obtained from the 3DIV database (available at http://3div.kr/), specifically the tissue mnemonics H1, ME, TB, MSC, NPC, LI11, PA, LG, and GM. The UK Biobank fine-mapped eQTL data were downloaded from the supplementary material of Wang et al.’s study^51^. The summary statistics were downloaded from the IEU GWAS portal^62^ (IDs: ieu-b-7, ebi-a-GCST90027158, ebi-a-GCST90027158, ebi-a-GCST005647, ebi-a-GCST90091033, ebi-a-GCST90091033, ebi-a-GCST90038627, ieu-b-5099, ieu-a-1185, ieu-b-5110) and for hepatitis, from the BioStudies database^63^ (ID: S-BSST407). All reference datasets used to run s-LDSC^65^ were downloaded following the links from the source material: https://github.com/bulik/ldsc. The model architectures and all training and analysis scripts, along with scripts to download and complete all pre-processing steps on the training data (sourced from Roadmap^16^ and largely replicated from Chromoformer’s scripts^17^) are available at https://github.com/neurogenomics/chromexpress.

