## Supplementary Material for "Predicting gene expression from histone marks using chromatin deep learning models depends on histone mark function, regulatory distance and cellular states"

| Epigenome ID (EID) | Shorthand name | GROUP | Epigenome Mnemonic | ANATOMY | TYPE |
|---|---|---|---|---|---|
| E118 | Hepatocellular carcinoma | ENCODE2012 | LIV.HEPG2.CNCR | LIVER | CellLine |
| E116 | Lymphoblastoid | ENCODE2012 | BLD.GM12878 | BLOOD | PrimaryCulture |
| E114 | Lung carcinoma | ENCODE2012 | LNG.A549.ETOH002.CNCR | LUNG | CellLine |
| E087 | Pancreatic islets | Other | PANC.ISLT | PANCREAS | PrimaryTissue |
| E066 | Liver | Other | LIV.ADLT | LIVER | PrimaryTissue |
| E016 | HUES64 ESC | ESC | ESC.HUES64 | ESC | PrimaryCulture |
| E007 | H1 derived NPC | ES-deriv | ESDR.H1.NEUR.PROG | ESC_DERIVED | ESCDerived |
| E006 | H1 derived MSC | ES-deriv | ESDR.H1.MSC | ESC_DERIVED | ESCDerived |
| E005 | H1 derived trophoblast | ES-deriv | ESDR.H1.BMP4.TROP | ESC_DERIVED | ESCDerived |
| E004 | H1 derived mesendoderm | ES-deriv | ESDR.H1.BMP4.MESO | ESC_DERIVED | ESCDerived |
| E003 | H1 ESC | ESC | ESC.H1 | ESC | PrimaryCulture |

**Supplementary Table 1 Roadmap information on cell types.** ID and metadata for cell types taken from Roadmap.

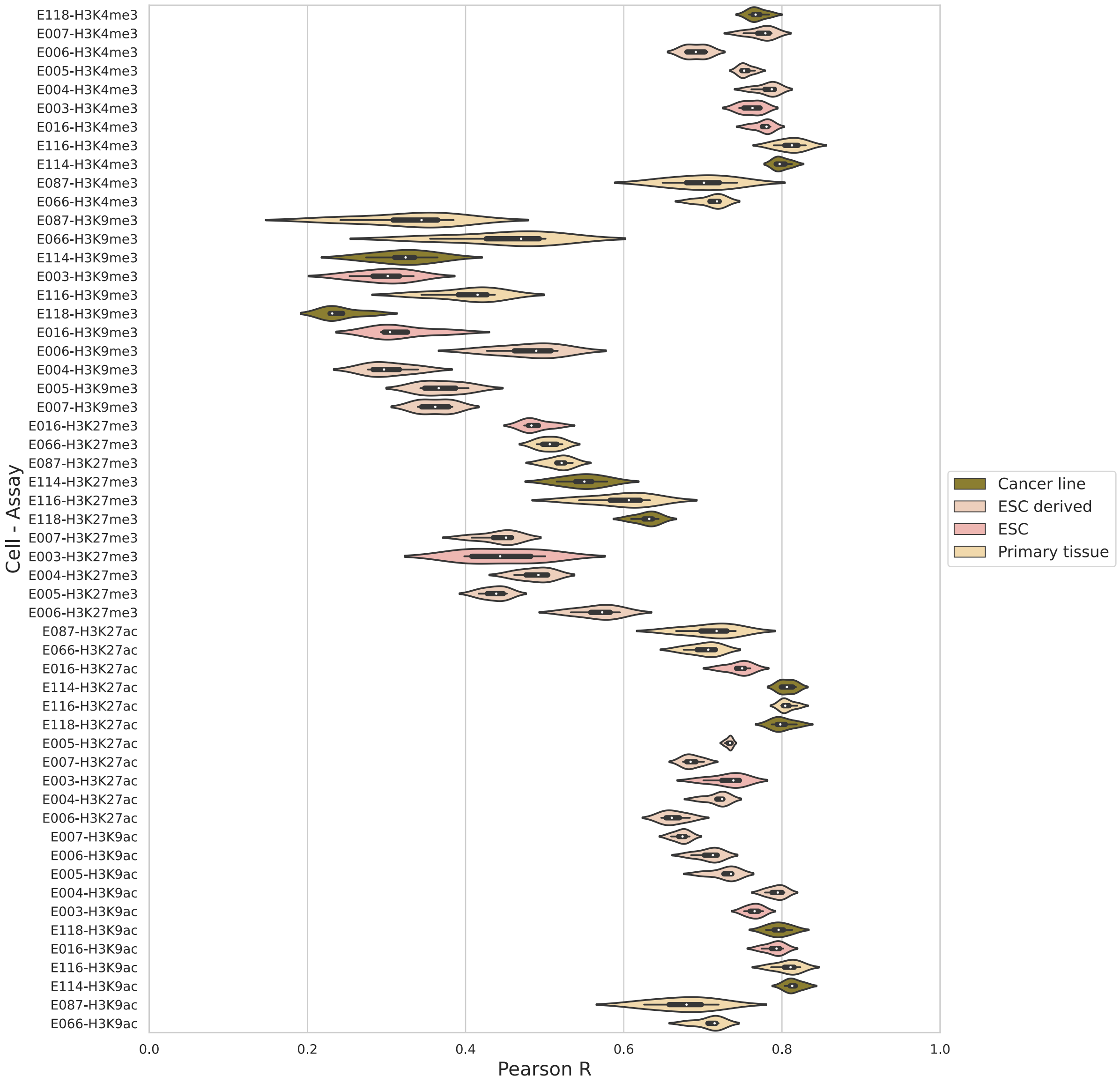


**Supplementary Fig. 1 Performance of promoter model by cell type and histone mark.** Violin plot of the model's performance on histone marks measured by Pearson correlation coefficient on the blind test sets. Here, the cell type performance is coloured by the cell state - Embryonic stem cell (ESC), ESC derived cell, adult primary tissue or cancer cell line. Splitting the performance by cell type and histone mark removes the multimodal distribution noted in **Fig. 2b**.

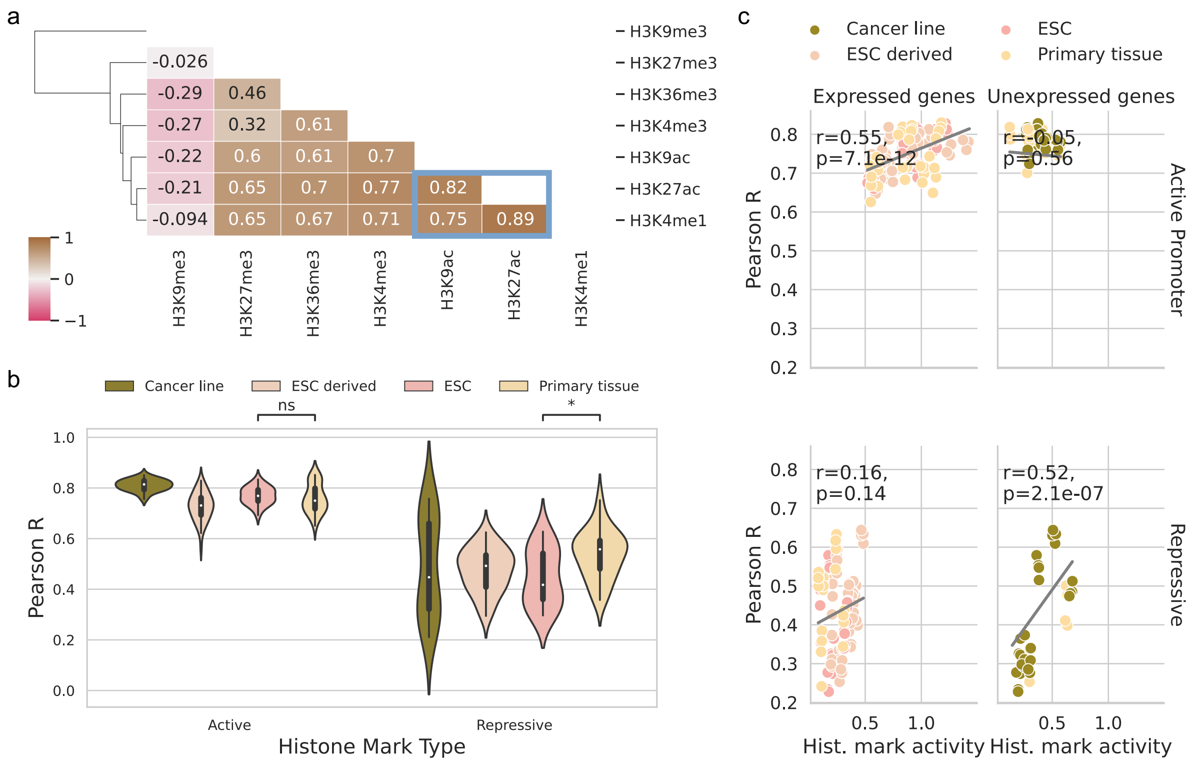


**Supplementary Fig. 2 Distal model's correlation of histone marks' function and performance by cell state.** (a) Correlation matrix of the distal model's performance by the different histone marks across each cell type and cross-validation k-fold. The distinct group of active marks from the promoter model, highlighted in blue, was also replicated for the distal model. Bars along the y-axis correspond to the hierarchical clustering dendrogram. (b) Violin plot of performance measured by Pearson correlation coefficient on the blind test sets grouped by cell state and histone mark levels. Significance based on Mann-Whitney U-Test with false discovery rate multiple test correction where p-value indicators: * < 0.05, ns >= 0.05. (c) Correlation between the histone mark levels (average log2-transformed, read depth in the full 40,000 base-pairs around the transcriptional start site) and model performance for each cell type, k-fold combination. Split by repressive and active marks and highly and lowly expressed genes.

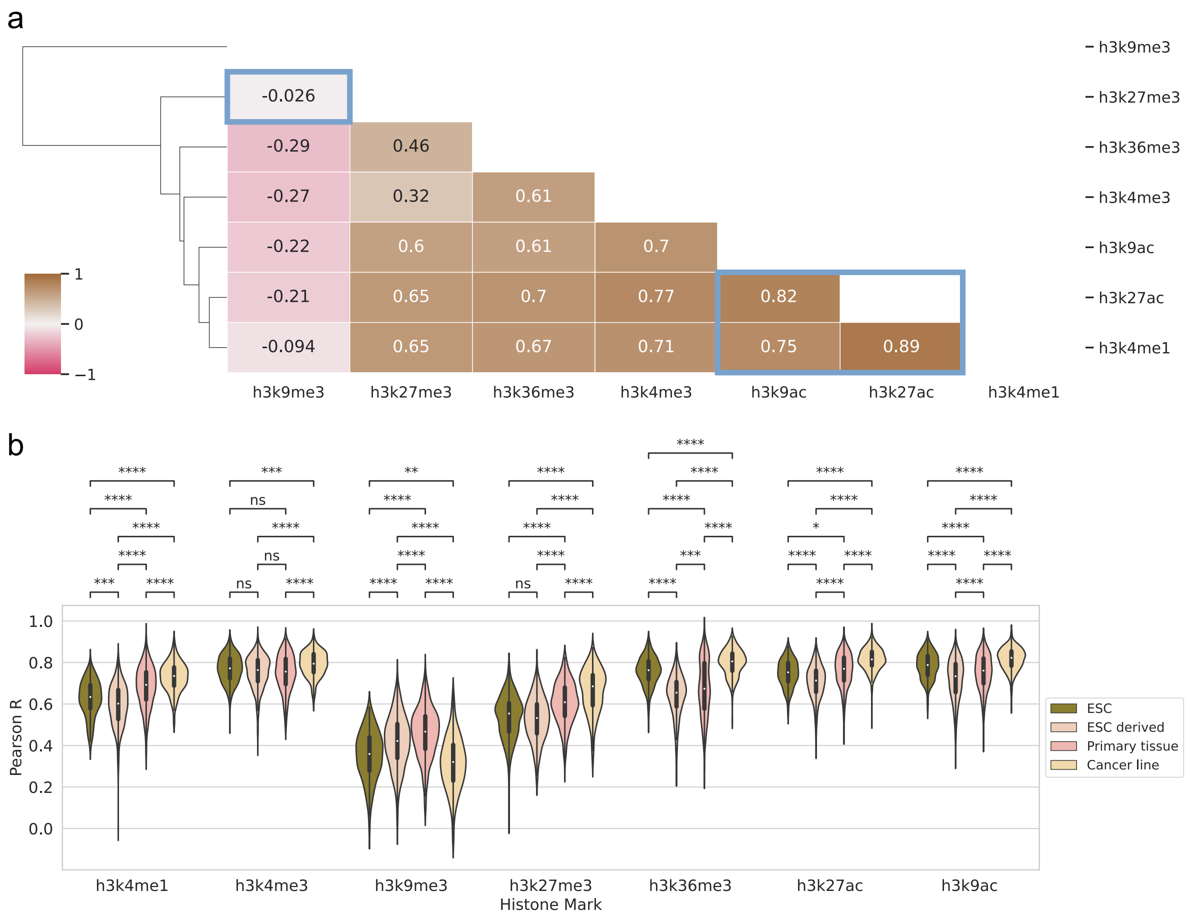


**Supplementary Fig. 3 Performance of distal model in gene expression predictions from two histone mark pairs with H3K27me3.** Performance for combinations of all pairs with H3K27me3, a repressive mark. The range of values represent the standard deviation across the different cell types and the 4-fold cross-validation. Data is averaged at the level of cell type and k-fold. The red dashed line shows the model's performance when trained on all seven histone marks together. Significance based on false discovery rate (FDR) multiple test correction where p-value indicators: **** < 1e-4, *** < 1e-3 and ** < 1e-2.

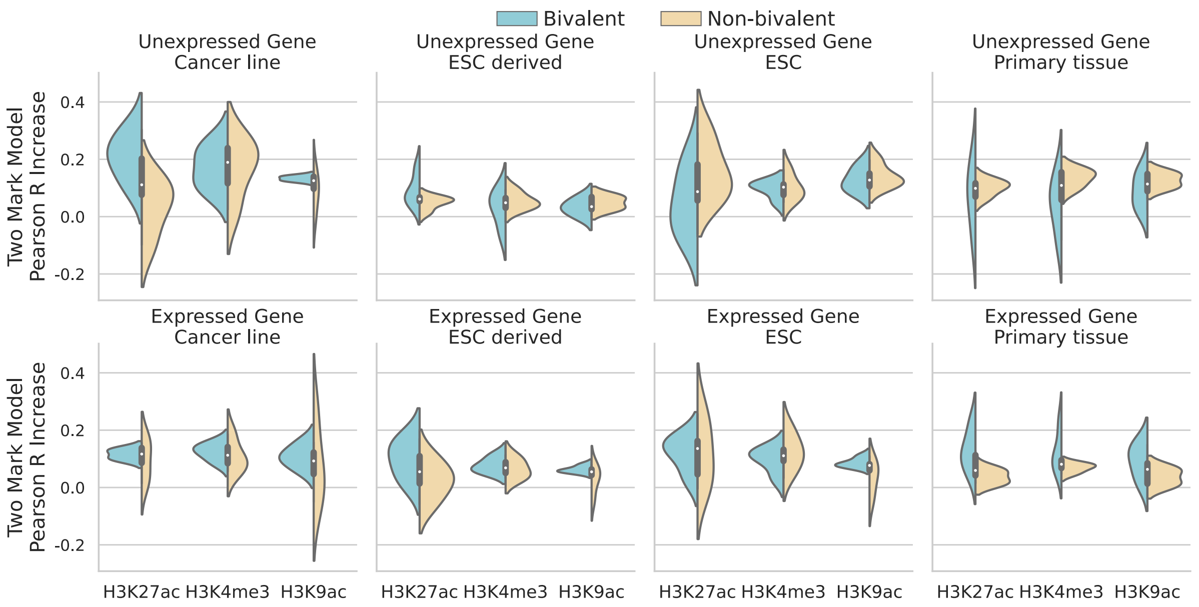


**Supplementary Fig. 4 Performance improvement of distal model from two histone mark pairs with H3K27me3 in bivalent genes.** Performance improvements for combination distal model over the single mark distal model for the top three, activating histone marks with H3K36me3. The range of values represent the standard deviation across the different cell types and the 4-fold cross-validation. Data is averaged at the level of cell type and k-fold. Performance improvement is split by mark, cell type group and bivalent and non-bivalent genes. Bivalent genes are defined as those with both an active histone mark signal and repressive - H3K36me3 signal in the gene promoter region (both above the median for the cell type). Plot is also split by highly and lowly expressed genes defined in the same way as for the histone mark level, above or below the median for the cell type.

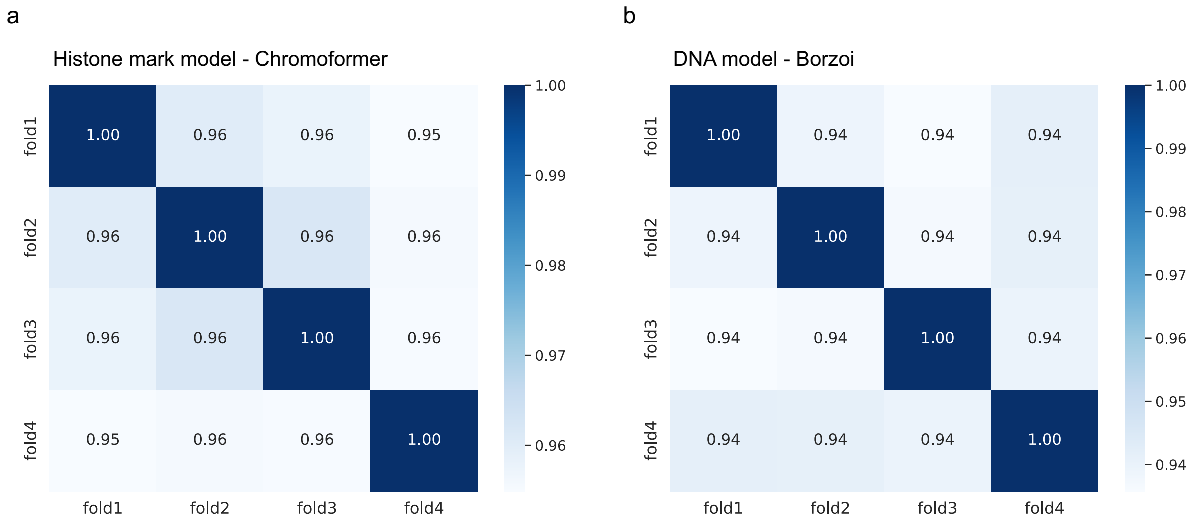


**Supplementary Fig. 5 Correlation across 4 fold cross-validation models in the *in silico* perturbation analysis.** (a) Correlation for Chromoformer, the histone mark model on matched genes and histone mark perturbation loci and level across the 4-fold, cross-validation models. All models showed high concordance (Pearson R>=0.95). (b) Correlation for Borzoi, a DNA model on matched genes and *in silico* mutagenesis levels across the 4-fold, cross-validation models. The same cell types and receptive field were inspected for correlation to chromoformer to make for a fair comparison. All models also showed high concordance (Pearson R=0.94) but slightly less than the histone mark model approach.

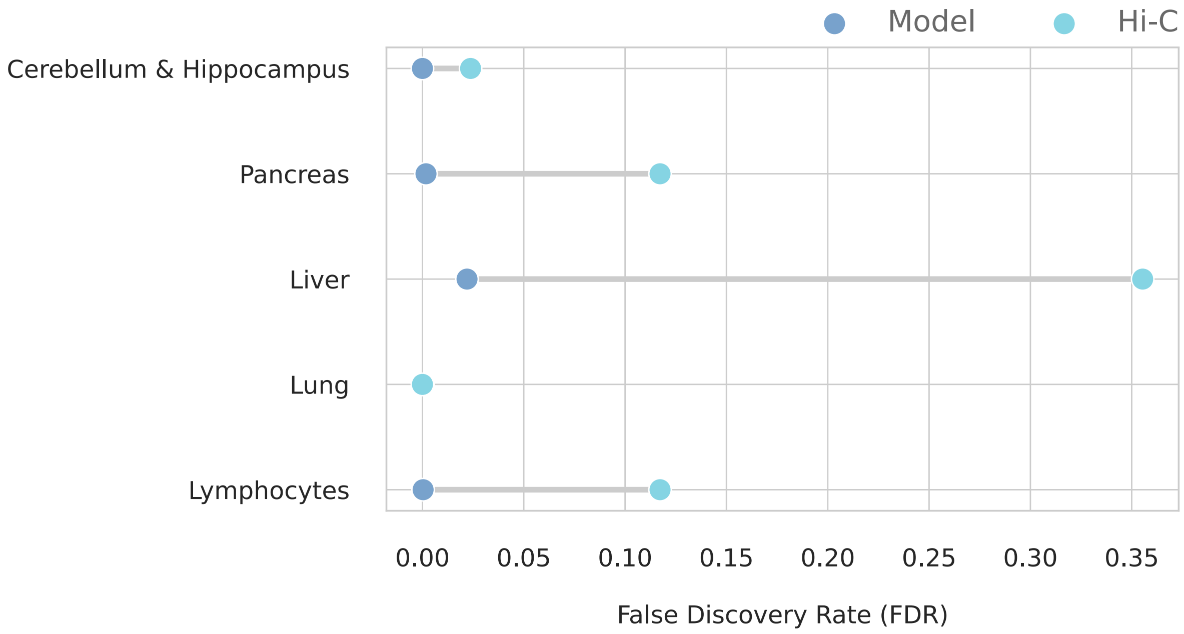


**Supplementary Fig. 6 Improvement of model over Hi-C data in fine-mapped eQTL enrichment.** Upstream *in silico* histone mark perturbation experiments from the active model were sorted into deciles based on their predicted change in expression and the top decile used (Model) to compare against cell type-specific Hi-C data upstream of matching genes (Hi-C). Both were tested for enrichment of fine-mapped eQTL interactions in matched cell types (y-axis) and compared against bootstrap sampling random upstream loci 10,000 times to generate p-values of enrichment (x-axis). Lower p-values for the model indicate greater enrichment found in more tests against randomly selected loci than the Hi-C data. For Lung, both the model and Hi-C data obtained the same p-value.
